# Meningeal lymphatic endothelial cells fulfill scavenger endothelial cell function and employ Mrc1a for cargo uptake

**DOI:** 10.1101/691477

**Authors:** Yvonne Padberg, Andreas van Impel, Max van Lessen, Jeroen Bussmann, Stefan Schulte-Merker

## Abstract

Brain lymphatic endothelial cells (BLECs) constitute a group of loosely connected endothelial cells within the meningeal layer of the zebrafish brain. We previously reported that BLECs efficiently endocytose extracellular cargo molecules (van Lessen et al., 2017), but how this is accomplished and controlled on the molecular level remains unclear. We here compare BLECs to scavenging endothelial cells (SECs) in the embryonic cardinal vein and find them to accept an identical set of substrate molecules. While there is redundancy in the type of scavenger receptors being used, the two cell populations rely for specific substrate molecules on different cell surface receptors to mediate their physiological role: Stab2 appears more critical within SECs in the cardinal vein, while BLECs depend more on the Mrc1a receptor for internalization of cargo. Given the striking similarities to the substrate specificity of cardinal vein SECs, we postulate that BLECs qualify functionally as SECs of the brain.

## Introduction

The lymphatic vascular system constitutes a blind-ended network that drains interstitial fluid and macromolecules from tissues and organs and eventually returns its contents back into the blood circulation. Furthermore, lymphatics are essential for immune cell trafficking and fat absorption in the intestine (Schulte-Merker et al., 2011). For a long time, the brain was considered to be devoid of lymphatic vessels, but recently a lymphatic vascular network in the dura mater of the mouse meninges was (re-)discovered (Mascagni and Bellini, 1816, Aspelund et al., 2015, Louveau et al., 2015). Lymphatic vessels are located in the immediate proximity of the meningeal blood vasculature where they drain cerebral interstitial fluid (ISF), macromolecules and cerebrospinal fluid (CSF) into deep cervical lymph nodes (Aspelund et al., 2015, Louveau et al., 2015). They develop postnatally, originating around the foramina that form the entry and exit sites for the blood vessels (BVs) and nerves, and migrate along blood vessels and the cranial and spinal nerves, eventually resulting in a fully developed lymphatic system at P28 (Antila et al., 2017). The meningeal lymphatic system appears to be conserved across mammals and has been described in humans and non-human primates (Absinta et al., 2017). Whether and how meningeal lymphatics might contribute to the waste removal of the brain has not been experimentally addressed but is a topic of significant interest, since accumulation of protein aggregates is one of the hallmark features of neurodegenerative diseases (Metcalf et al., 2012).

Recently we and others have demonstrated the presence of lymphatic endothelial cells in the meningeal layer of the zebrafish brain (Bower et al., 2017, Venero Galanternik et al., 2017, van Lessen et al., 2017). Due to the simultaneous but independent discovery of these cells, they were termed either brain lymphatic endothelial cells (BLECs) (van Lessen et al., 2017), mural lymphatic endothelial cell (Bower et al., 2017), or fluorescent granular perithelial (FGP) cells (Venero Galanternik et al., 2017) - all terms referring to the same cell type. These cells express lymphatic marker genes such as *prox1a, lyve-1* and *flt4 (vegfr3)*, develop in a Vegfc, Flt4 and Ccbe1-dependent manner and populate the menigeal layer of the brain. Even though BLECs are in close proximity to the meningeal blood vasculature, they do not share – in contrast to pericytes - a common basement membrane with the endothelial cells (van Lessen et al., 2017). Whole transcriptome profiling of sorted BLECs confirmed that BLECs are a distinct endothelial cell population, which show expression profiles different from macrophages and pericytes, while expressing lymphatic markers such as *flt4, lyve-1, prox1a* (Bower et al., 2017, van Lessen et al., 2017). In addition, inhibition of myelopoesis by administration of a pu.1 (spi1b) morpholino does not affect BLEC development, demonstrating that these cells do not constitute a macrophage lineage. Remarkably these cells do not form any vascular structure, but give rise to a network consisting of individual lymphatic endothelial cells expanding over the whole brain surface (van Lessen et al., 2017, Venero Galanternik et al., 2017, Bower et al., 2017).

BLECs originate from the venous choroidal vascular plexus behind the eye and sprout around 56hpf, at which point they downregulate blood vascular specific genes and upregulate lymphatic markers such as *flt4*. Sprouting occurs bilaterally and cells migrate along the mesencephalic vein (MsV), resulting in symetric loops of single cells that cover the optic tectum (TeO) of the zebrafish embryo. This network of individual cells subsequently expands throughout the development of the fish and covers the whole surface of the brain at around 3 weeks of age. BLECs have been shown to play a role in regenerative processes within the brain (Bower et al., 2017, Chen et al., 2019), and have an enormous ability to take up extracellular substances into subcellular vesicles in a process depending on receptor-mediated endocytosis (RME) (van Lessen et al., 2017). Previously we have shown that the uptake of avidin coupled to pHrhodo which is a pH-sensitive tag that only fluoresces upon internalization into the acidic compartment of the lysosome, can be blocked by mannan. This suggests that the mannose receptor is involved in the uptake of avidin. In line with the high endocytotic capacity of these cells, which becomes evident immediately upon sprouting from the choroidal vascular plexus, BLECs typically have large spherical vacuoles which are interpreted as lysosomal compartments in adult brains (van Lessen et al., 2017, Venero Galanternik et al., 2017).

Another endothelial subpopulation, that has been reported to posses a high endocytotic capacity, are scavenger endothelial cells (SECs). In all terrestrial vertebrates this specialized cell population is located in the liver sinusoids and termed liver sinusoidal endothelial cells. In teleost fish, sharks and lampreys it was identified in various other organs (Seternes et al., 2002). In embryonic zebrafish, SECs were recently shown to be present in several large veins, including the posterior and common cardinal vein (PCV, CCV), and the caudal vein (CV) where they clear substances, colloidal waste and viral particles from the blood circulation as early as 28hpf. The uptake of cargo molecules by SECs in the CV is mainly dependent on the transmembraine receptor *stabilin-2* (Campbell et al., 2018). The questions arise whether BLECs possibly represent a functional equivalent of this cell population serving scavenger functions of the brain and whether they might substitute for the absence of lymphatic vessels in teleost meninges.

Here we explore this hypothesis and compare BLECs with SECs in the CV to study the physiological role of BLECs. We have generated and analyzed mutants for some of the classical cargo receptor molecules and tested whether they are required for the internalization of cargo into BLECs. We showed that BLECs are highly endocytic cells, which are are considerably more efficient in macromolecular uptake for the tested substrate than microglia. BLECs and SECs in the CV have the same substrate specificity and take up a range of macromolecules including proteins, liposomes, lipoproteins, polysaccharides and glycoaminglycans. Due to the efficiency of cargo internalization and the similarities of the type of cargo they are able to endocytose, we conclude that BLECs qualify as a novel population of-scavenger endothelial cells residing in the brain area. Surprisingly, we find that both cell populations depend on different receptors mediating endocytosis.

## Results

### BLECs and SECs within the caudal vein share the same substrate specificity

In order to investigate the function of BLECs in the zebrafish embryo, we directly compared SECs in the CV with BLECs. To this end we injected the same macromolecules either into the optic tectum of embryos (Figure 1 A-J) or into the blood circulation (Figure 1 K-U) at 5dpf to compare the substrate specificity of both cell populations. In most cases at least two dyes were co-injected and the intracellular uptake was monitored. We used different classes of dye-conjugated substrate molecules, including liposomes (DOPG liposomes - Figure 1D, U), modified lipoproteins (oxidated-LDL - 1F, Q), glycoaminoglycans (not shown), proteins such as Avidin (Figure 1 G, R), Transferrin (Figure 1H, S) and Amyloid-β (Figure 1 J, P) and the polysaccharide dextran (Figure 1 E,T). Without exception, these molecules or particles were taken up by both cell populations, suggesting that they share the same substrate specificity (Figure 1V), even though the two cell populations serve two completely different anatomical and physiological compartments: BLECs clear the ventricles and the brain extracellular space from macromolecules, while SECs in the CV filter out substrates from the blood stream.

**Figure 1:**
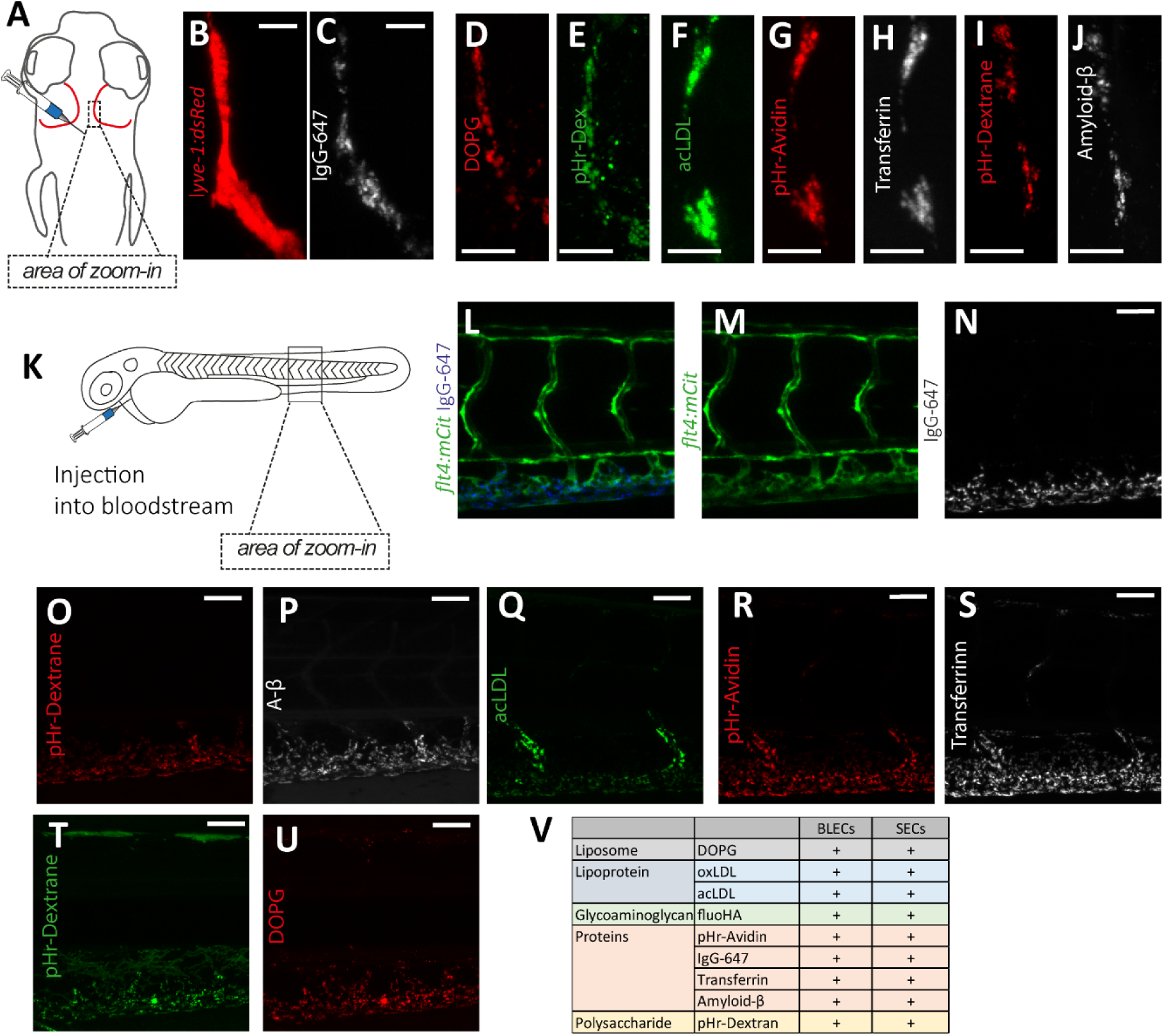
Brain lymphatic endothelial cells (BLECs) in the meningeal layer and sinusoidal endothelial cells (SECs) in the common cardinal vein (CCV) share the same substrate specificity. (A) Overview of the zebrafish head region (dorsal view, anterior at the top) with the position of BLECs highlighted in red. The boxed area indicates the position of the imaged BLECs shown in B-J. Injection of different fluorescent substrate molecules was performed either into the center of the optic tectum (TeO), close to the meninges, or into the cerebrospinal canal, imaged in C-J. (B,C) Uptake of IgG-Alexa647 by BLECs. Confocal projections of a *lyve1:DsRed* positive BLEC (B), which has internalized the fluorescent IgG-Alexa647 (C) that was administered as described in A. (D-J) Confocal projections of different classes of DOPG (D), dextran-488 (E), acLDL (F), pHr-Avidin (G), Transferrin (H), dextran-565 (I) and Amyloid-β (J) internalized by BLECs. A-J Scale bar represents 12.5μm. (K) Schematic overview of a zebrafish embryo, indicating injections of different fluorescent dyes into the blood stream. The boxed area highlights the location of the CCV that was imaged in (L-U). (L) Confocal pictures of *flt4:mCitrine* transgene (M) and the uptake of IgG-Alexa647 (N), pHr-Dextran-565(O), Amyloid-β (P), acLDL (Q), pHr-Avidin (R), Transferrin (S), pHr-Dextran-488(T) and DOPG (U) by SECs located in the CV. (L,M,N,T,U) *mrc1a* ^*+/*−,^ (O-S) wild type embryo. (V) Summary of the fluorescent substrate molecule uptake experiments; K-U scale bar represents 50μm. pHr-Dex – pHr-Dextrane; DOPG - 1,2-dioleoyl-sn-glycero-3-phospho-(1′-rac-glycerol); acLDL-acetylated LDL; oxLDL – oxidated LDL; fluoHA-fluorescent hyaluronic acid; IgG-647 – IgG-Alexa647

### BLECs are more efficient in tracer uptake than macrophages

A major open question is which function BLECs fulfill on the surface of the brain. Our initial results suggested a role in clearance of extracellular waste products. However, microglia have always been considered as the major force to remove cellular or sub-cellular components from the brain (Platt et al., 1998). We therefore asked how the endocytic capacity of microglia compares to that of BLECs. We injected IgG-Alexa647 into *lyve-1:dsRed;mpeg:GFP* double transgenic embryos that allow the discrimination of BLECs and microglia (Figure 2A). We confirmed that IgG-Alexa647 was taken up by nearly all of the *mpeg:GFP*^*+*^ microglia (Figure 2B). Surprisingly, the fluorescence of IgG-Alexa647 that accumulated in the vacuoles of the microglia was significantly lower in intensity than that of IgG-Alexa647 accumulating in the BLECs (Figure 2C red/white arrow heads). In order to quantify this phenomenon and to analyze the dynamics of the dye uptake, IgG-Alexa647 was injected into 5dpf embryos and the fluorescence intensity of IgG-Alexa647 signal in BLECs was quantified (Region 1 and 2 depicted in Figure D-D’’ and E) and compared to the intensity of the region between the 2 loops (region 3, Figure D-D’’ and E), where *mpeg:GFP* positive microglia are located, at 1 hour post injection (hpi), 3hpi and 6hpi. At all three time points, BLECs were found to take up significantly more IgG-Alexa647 than *mpeg* positive microglia (Figure 2H, n=6 1hpi** p< 0,005; 3hpi **p<0,005; 6hpi **p<0,005). Hence, while microglia are in closer proximity to the ventricles in which the dye has been injected, BLECs were still an order of magnitude more efficient in the internalization of IgG-Alexa647.

**Figure 2:**
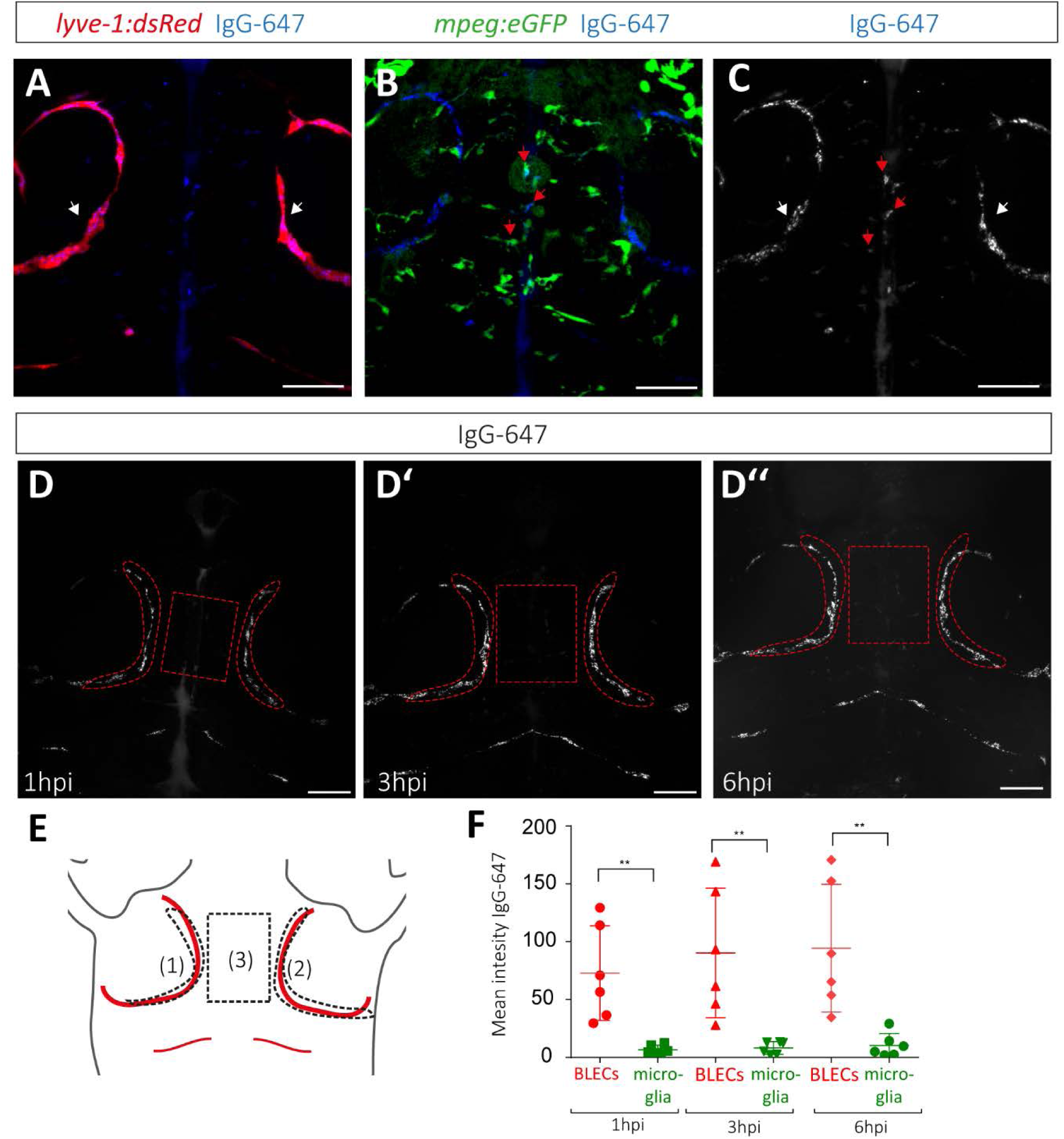
BLECs on the optic tectum are more efficient in the uptake of exogenous substrate than microglia. (A-C) Confocal projection of the head region (dorsal view, anterior to the top) of a *lyve-1:dsRed;mpeg:eGFP* double transgenic embryo, injected with IgG-Alexa647 at 5dpf. (A) Composite of *lyve1* expressing BLECs (white arrow) and IgG-Alexa647. (B) Composite of *mpeg:eGFP* positive microglia and IgG-Alexa647. As an example, three IgG-Alexa647 accumulating microglia cells are highlighted with red arrows. (C) Maximum projection of IgG-Alexa647. (D’-D’’) Maximum projection of the same IgG-Alexa647 injected embryo at 1hpi (D), 3hpi (D’), and 6hpi (D’’). (E) Schematic overview of the regions that were used for the uptake quantifications: BLECs within the left loop (1), the right loop (2), and the microglia positioned in between the loops (3). The identical areas are also indicated in the confocal projections in D-D’’. (F) Quantification of the mean intensity above zero of IgG-Alexa647 of the two loops (region1+region2)/2 versus microglia cells at 1hpi (test p=0,0027), 3hpi (Mann-Whitney test p=0,0022) and 6hpi (Mann-Whitney test p=0,0022). The scale bar indicates 50μm. ** equals p < 0,01. BLECs-brain lymphatic endothelial cells; IgG-647 – IgG-Alexa647

### *mrc1a* mutant zebrafish show an increase in BLEC numbers

The mannose receptor Mrc1a has previously been shown to be highly expressed in BLECs and is suggested to mediate endocytosis of at least one substrate - avidin - via receptor-mediated endocytosis (van Lessen et al., 2017). In mice, the mannose receptor is maintaining homeostasis of macromolecules in the blood and is responsible for the uptake of Lutrophin (Mi et al., 2002), denaturated collagens (Malovic et al., 2007) and serum glycoproteins including most lysosomal hydrolases (Lee et al., 2002). We have previously shown that mannan, which is a bacterial polysaccharide binding very efficiently to the mannose receptor (Sallusto et al., 1995), could block the uptake of pHr coupled Avidin by BLECs (van Lessen et al., 2017). However, other mannan-binding proteins such as mannan-binding lectin exist. In order to investigate whether zebrafish Mrc1a is mediating the removal of macromolecules from the brain and acts as a clearance receptor, we generated a mutant for *mrc1a* harbouring a frameshift mutation within exon4 (Figure 3A). Homozygous mutants did not show any obvious lymphatic or blood vessel phenotype at 5dpf (Figure 3 B,C), and were identified in a normal Mendelian ratio as viable and fertile adults.

**Figure 3:**
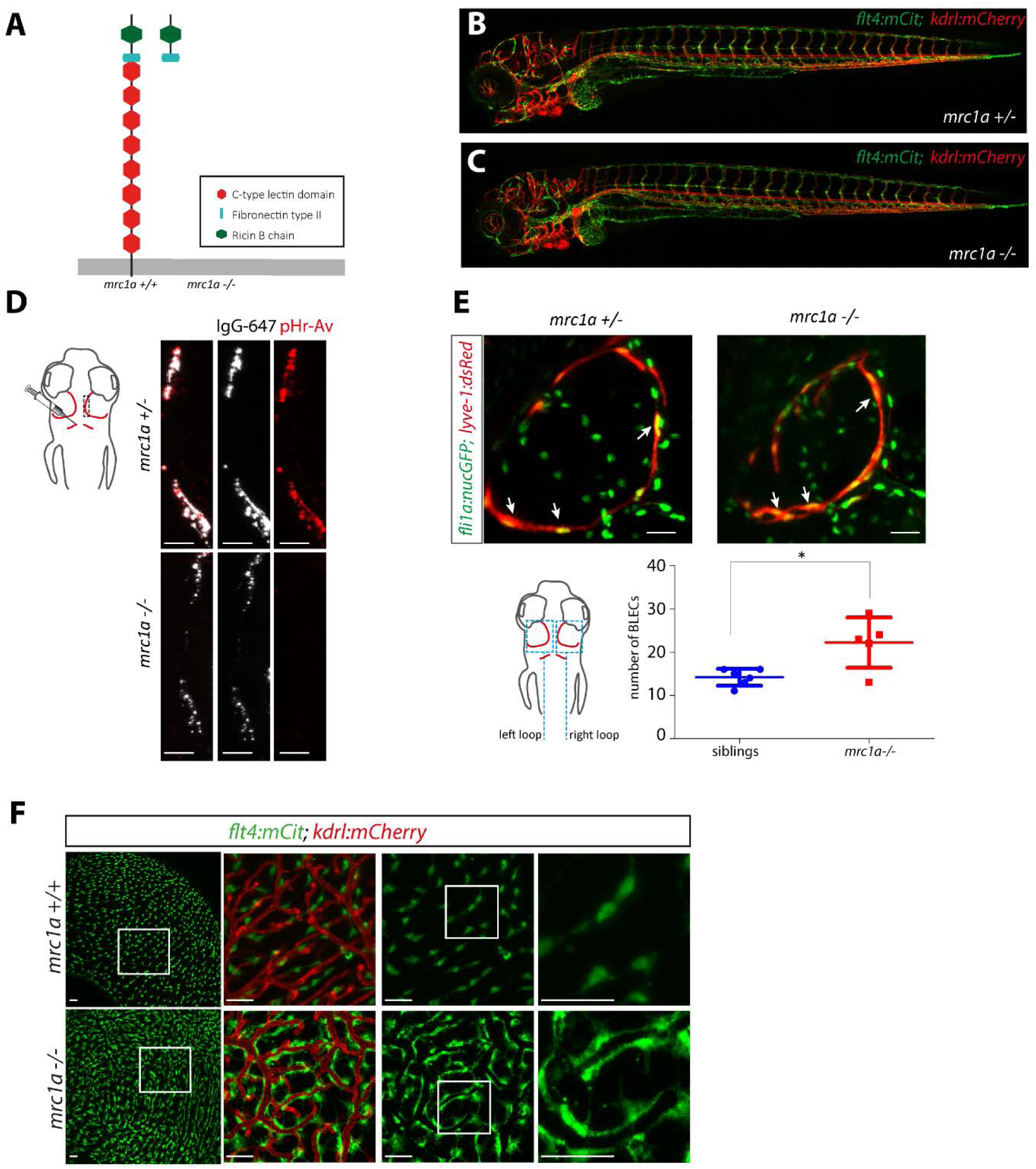
*mrc1a* is indispensable for the uptake of pHR-Avidin. (A) Predicted wild type Mrc1a domain structure and predicted protein structure of the mutant allele. (B,C) Confocal picuture of *flt4:mCit; kdrl:mCherry* double transgenic embryos at 5dpf, with no obvious lymphatic or vascular phenotype defects in *mrc1a* homozygous embryos. (D) Confocal picture of embryos injected with IgG-Alexa647 and pHR-Avidin, demonstrating that the uptake of pHR-Avidin is blocked in *mrc1a* mutant embryos, whereas the uptake of IgG-Alexa647 is not affected. Scale bar represents 12,5μm. (E) Confocal picture of *lyve1a:dsRed;nucfli1a:GFP* double transgenic embryos. BLEC numbers in both loops on the optic tectum were counted in *Tglyve1a:dsRed;nucfli1a:GFP* embryos. 3 arrows are pointing at 3 positive cells, co-expressing both lyve1a and nucfli1a:GFP. Scale bar represents 25μm. *mrc1a* mutants have on average 22 BLECs in both loops (n=5) whereas their siblings have 14 BLECs (n=6) at 5dpf (mean ±SEM measured in both loops) (t-test sibling versus *mrc1a*^−*/*−^* p=0,012). (F) Brains of 4 months old adult fish Tg(*flt4:mCit;kdrl:mCherry*) highlighting BLECs (green) and blood vessels (red). Squares indicate the enlarged regions to the immediate right. The scale bar represents 50μm. pHr-Av – pHr-Avidin

Interestingly, upon quantification of the number of BLECs in both loops of the optic tectum of the 5dpf zebrafish brain in double transgenic embryos *lyve-1:dsRed;fli1a:nucGFP*, we found a highly significant increase (56%) in BLEC numbers in *mrc1a* mutants compared to sibling controls (Figure 3E). The difference in cell numbers between heterozygous and homozygous *mrc1a* embryos was already visible at 4dpf (data not shown). This increase in BLEC cell number was also evident in whole brains of adult *mrc1a* mutant fish. In addition, we found that BLECs display an altered morphology in adult *mrc1a* mutants (Figure 3F).

### *mrc1a* mutants are deficient in pHr-Avidin uptake

When IgG-Alexa647 and pHr-Avidin was co-injected into an *mrc1a*^*+/*−^ incross, we found that siblings show an uptake of both dyes into BLECs, whereas *mrc1a* mutant embryos only endocytosed IgG-Alexa647 but did not take up pHR-Avidin (Figure 3D). We repeated the experiments co-injection of mannan with IgG-Alexa647 (van Lessen et al., 2017) and found that IgG-Alexa647 was not only retained at the plasma membrane level, but was completely endocytosed into BLECs whereas pHr-Avidin uptake was blocked completely after mannan administration. When we injected pHr-Avidin first, followed by mannan and IgG-Alexa647, the two dyes were indeed found in identical lysosomes (Supplement S1). We conclude that competitive inhibition using mannan phenocopies the *mrc1a* mutant phenotype. Analysis of *mrc1a* mutants demonstrated that the endocytotic uptake of pHr-Avidin, but not of IgG-Alexa647 by BLECs depends on the Mrc1a receptor.

### BLECs and SECs in the CV have different molecular mechanisms for dye internalization

Stabilin-2 (Stab2) has recently been found to represent an essential receptor for SEC function in the zebrafish caudal vein, clearing the circulation from nanoparticles such as fluorescently labelled hyaluronic acid (fluoHA) and liposomes (Campbell et al., 2018) (Figure 4A). We therefore asked whether Mrc1a also mediates macromolecule uptake within the caudal vein and whether there is redundancy between Mrc1a and Stab2.

**Figure 4:**
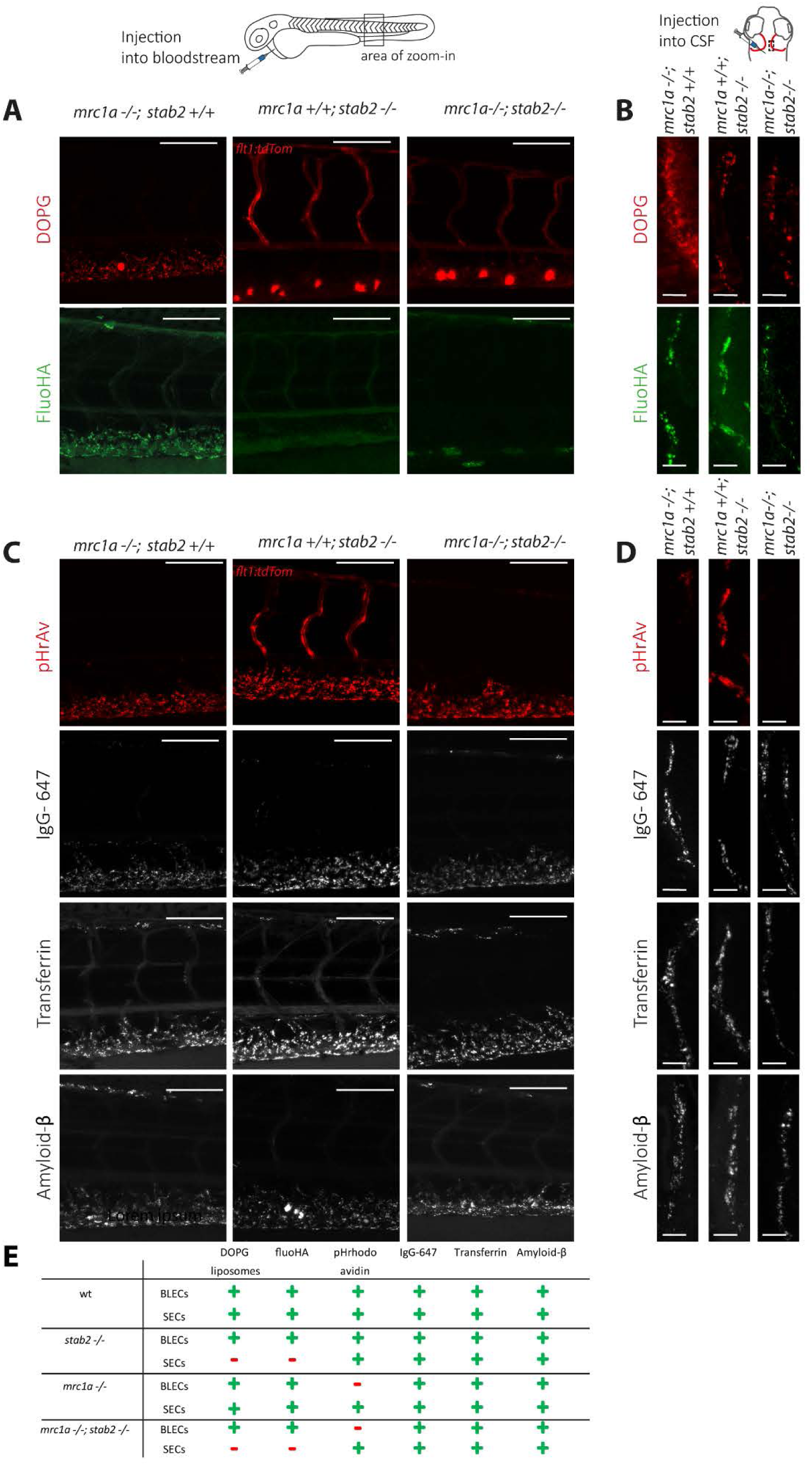
Differential involvement of Stab2 and Mrc1a receptors during the uptake of cargo molecules by BLECs and SECs. Analysis of dye uptake into SECs and BLECs of either *mrc1a* ^−*/*−^ or *stab2*^−*/*−^ single mutants, or of *mrc1a*^−*/*−^; *stab2*^−*/*−^ double mutant embryos. (A) Confocal images showing the accumulation of DOPG and fluoHA within SECs of the CV after dye administration into the bloodstream of embryos with the indicated genotypes. (B) DOPG and fluoHA uptake by BLECs after injection into the CSF or the optic tectum. (C) Confocal picture showing uptake of pHr-Avidin, IgG-Alexa647, Transferrin and Amyloid-β by SECs, which were administered to the bloodstream. (D) Confocal projections of pHr-Avidin, IgG-Alexa647, Transferrin and Amyloid-β uptake by BLECs, after injection into the CSF or the optic tectum. (E) Table summarizing the uptake of the different dyes by BLECs and SECs. Scale bar in A,C represents 100μm, and in B,D 12.5μm, respectively. pHr-Av – pHr-Avidin; DOPG - 1,2-dioleoyl-sn-glycero-3-phospho-(1′-rac-glycerol); fluoHA-fluorescent hyaluronic acid; IgG-647 – IgG-Alexa647

To answer these questions, we generated *mrc1a*^−*/*−^;*stab2*^−*/*−^ double mutants and compared substrate specificity in wt, *mrc1a*^−*/*−^, *stab2*^−*/*−^ and double mutants. We found that the internalization of two known substrates for SEC clearance in the caudal vein - hyaluronic acid and DOPG-liposomes-were affected only in *stab2* single and *mrc1a*^−*/*−^;*stab2*^−*/*−^ double mutants. Single *mrc1a* mutants, however, did not exhibit any defects, indicating that Mrc1a is not essential for clearance of these ligands (Figure 4 A). Next we wanted to assess the importance of *mrc1a* and *stab2* for the uptake of hyaluronic acid and DOPG liposomes in BLECs. Interestingly, neither *stab2*^−*/*−^ or *mrc1a*^−*/*−^ single, nor *mrc1a*^−*/*−^;*stab2*^−*/*−^ double mutants showed a defect in endocytosis of DOPG liposomes or hyaluronic acid in BLECs (Figure 4B) suggesting that other receptors must be involved in the endocytosis of these substrates. These results indicate, that even though SECs in the CV and BLECs have identical substrate specificity, there are important differences in the clearance mechanisms between the two cell types.

Since zebrafish Mrc1a is essential for the internalization of pHr-Avidin in BLECs, we wondered whether the Mrc1a receptor is mediating preferentially protein uptake in either BLECs or caudal vein SECs. We therefore tested the uptake of additional proteins such as pHr-Avidin, IgG-Alexa647, transferrin and amyloid-β in both cell types in the different mutants. Interestingly, neither Mrc1a nor Stab2 are essential for protein uptake by the SECs of the caudal vein from the blood plasma (Figure 4C). Similarly, when we analyzed the BLECs, we found that nearly all proteins were still cleared from the brain in *mrc1a, stab2* and double mutant embryos. Also the endocytosis of modified lipoproteins (acetylated and oxidized LDL) and dextran by BLECs and SECs was unaffected (Supplemental Figure 2). The only exception was pHr-Avidin, whose uptake by BLECs is completely blocked in the *mrc1a*^−*/*−^ single and *mrc1a*^−*/*−^;*stab2*^−*/*−^ double mutants (Figure 4D, E).

### *mrc1a*^−*/*−^ BLECs are less efficient in the uptake of dextran and IgG-Alexa647

Although all tested molecules (except for pHr-Avidin) and particles were still endocytosed by BLECs in *mrc1a* mutants, we noticed differences in the amount of endocytosed material between the different genotypes in some instances. To quantify this effect, we co-injected acLDL-488, pHr-Dextran and IgG-Alexa647 into *mrc1a* mutants and siblings and subsequently measured the fluorescence intensity of the dye accumulating in the vesicles of the BLECs as depicted in Figure 5A and B. Importantly, this analysis demonstrated that the amount of pHr-Dextran and IgG-Alexa647 accumulating in the vesicles of *mrc1a* mutant BLECs was significantly reduced compared to wild type sibling controls (t-test, **** p<0.0001). The amount of endocytosed acLDL was unaltered in *mrc1a* mutants. This shows that even though the *mrc1a* mutant fish can still endocytose pHr-Dextran and IgG-Alexa647 in the absence of a functional Mrc1a receptor, dye internalization is much less efficient suggesting that the uptake of dextran and IgG-Alexa647 are partially mediated by *mrc1a.*

**Figure 5:**
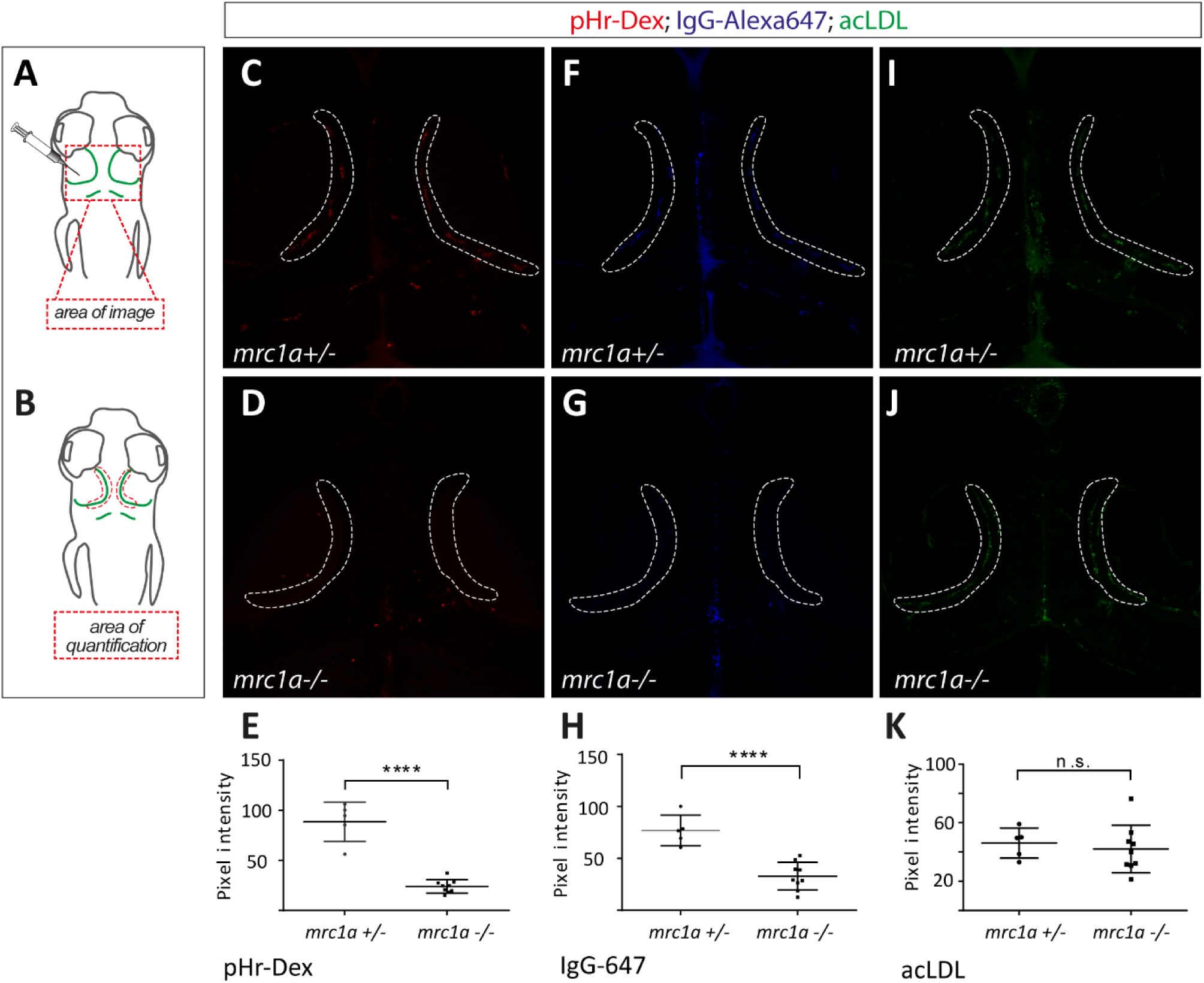
*mrc1a* mutants show a significantly reduced uptake of dextran and IgG-647 by BLECs. (A) Cartoons depicting the imaging area within the head region and the injection site of the different fluorescently labelled molecules. Dorsal view and anterior to the top in all images. All three substrates (acLDL-488; pHR-Dextran-564 and IgG-Alexa647) were co-injected into the same embryo. A total of 9 mutants and 6 heterozygous embryos were analyzed for each dye. (B) Overview of the area used for quantification, which is also indicated in the confocal projections (C,D,F,G,I,J). (C,D,E) *mrc1a* mutant BLECs take up significantly less dextran (mean ±SEM measured in each loop) (t-test sibling versus *mrc1a*^− */*−^ **** p<0.0001) and less IgG-647 (F,G,H) (mean ±SEM measured in each loop) (t-test sibling versus *mrc1a*^−*/*−^ **** p<0.0001) compared to their heterozygous siblings. (I, J, K) No difference in uptake by the BLECs was found in case of acLDL (mean ±SEM measured in each loop) (t-test sibling versus *mrc1a*^−*/*−^, p=0,63). Pixel intensity was analyzed using Fiji software. pHr-Dex – pHr-Dextrane; acLDL-acetylated LDL; IgG-647 – IgG-Alexa647

Taken together, our results demonstrate that BLECs are highly endocytic cells that accept the same substrate molecules as SECs in the CV and qualify as scavenger endothelial cells of the brain. However, the two cell populations rely at least in parts on different receptor molecules for cargo internalization. Whereas Mrc1a is essential for endocytosis in BLECs and thereby likely plays an essential role in waste removal from the brain, it is not crucial for waste removal from the blood by SECs in the CV. On the other hand, Stab2 is essential for internalization of selected substrates by the caudal vein SECs, but appears dispensable for BLECs.

## Discussion

Clearance of macromolecules from the brain parenchyma is a crucial process and has always been considered to be carried out by microglia in vertebrates. Recently, teleost BLECs were discovered, which cover the meningeal layer of the optic tectum and other parts of the brain as an extensive network of loosly connected cells in close proximity to blood vessels. They remain as single cells and never form lumenized structures, even after having expanded significantly during growth of the individuum. The question which function these cells fulfill under normal physiological conditions has not been answered. In response to cerebrovascular damage, however, BLECs are able to popupulate the brain parenchyma and can fulfill a guidance function for regrowing blood vessels (Chen et al., 2019).

It is striking that within the same anatomical compartment (meninges) of teleosts and mammals there are lymphatic endothelial cells to be found, but that these cells form different structures while still possibly serving similar physiological functions such as waste removal. Since BLECs cannot, in the absence of lumenized vessels, mechanistically work the same way as mammalian lymphatics, we here investigated whether BLECs represent a scavenger endothelial cell population of the brain. We compared BLECs to a recently discovered SECs in the CV of the zebrafish embryo, which has been demonstrated to clear the blood from macromolecules (Campbell et al., 2018), and in that sense are functionally homologous to mammalian liver sinusoidal endothelial cells. While trying numerous different substrate classes such as proteins, liposomes, lipoproteins, polysaccharides and glycoaminoglycans, we could not find any difference in the substrate specificity of caudal vein SECs and BLECs. We therefore conclude that BLECs function as *bona fide* scavenger endothelial cells.

Since microglia were always considered to constitute a cleaning mechansim which responds efficiently to tissue damage and infection by engulfing and processing pathogens (Platt et al., 1998), we studied the functional capacity of BLECs and microglia in direct comparison and analyzed how BLECs behave compared to macrophages in terms of internalization of cargo. Apart from their single cellular morphology and the expression of the mannose receptor, they have little in common. First, microglia mediate waste removal mainly via phagocytosis (Kettenmann, 2007, Barron, 1995), whereas BLECs take up their cargos via endocytosis (van Lessen et al., 2017). Second, BLECs are venous derived cells and sprout from the choroidal vascular plexus in the head (van Lessen et al., 2017, Venero Galanternik et al., 2017, Bower et al., 2017) whereas microglia stem from a hematopoietic precursor population (Goldmann et al., 2016). Third, BLECs and microglia are very different in motility. BLECs form a network of stationary cells, whereas microglia are distributed throughout the brain and use their long cellular branch extensions to remove dying neurons from the extracellular space (Mazaheri et al., 2014). The latter would suggest that microglia might be more efficient in clearing the brain from macromolecules, particularly since they reside in the brain parenchyma while BLECs are not in direct contact with neural tissue. By directly comparing microglia and BLECs efficacy, we found that BLECs are significantly more efficient in clearing the brain from the protein we tested, compared to macrophages at 1hpi, 4hpi and 6hpi (Figure 2H). This is counterintuitive, for the reasons mentioned above. Hence, we conclude here that BLECs fulfill the function of scavenger endothelial cells of the brain that can internalize IgG-Alexa647 in a very efficient way. Interestingly, within the mammalian liver, there is a similar division of labour between liver sinusoidal endothelial cells and the liver resident macrophages (Kupffer cells), which together form the reticuloendothelial system (RES) within this organ, clearing the blood plasma from endogenous and exogenous waste. We suggest that BLECs and microglia possibly function in a smilar way in the brain.

Concerning the spectrum of possible scavenger receptor molecules, we have focused on Mrc1a and Stab2. We found that Mrc1a mediates the uptake of several substrates (Figure 2D, Figure 5E,H) in BLECs. Avidin is so far the only substrate that we identified, that completely failed be taken up by the *mrc1a* mutant cells. Importantly, avidin that is isolated from hen egg white contains abundant high-mannose glycans (Fiete et al., 1997, DeLange, 1970, Green and Toms, 1970, Bruch and White, 1982). Glycoproteins containing these glycans are known ligands for the mannose receptor and of such substances are rapidly cleared from the blood circulation via liver sinusoidal endothelial cells in mammals (Hubbard et al., 1979). In contrast, transferrin normally does not contain abundant high-mannose glycans, providing an explanation for the selective requirement for Mrc1a in avidin clearance.

Although the other substrates we injected can be internalzed via alternative pathways, we show that also the pHr-Dextran and IgG-Alexa647 uptake is significantly reduced in *mrc1a* receptor mutants, highlighting the key role of Mrc1a in the removal of different substance classes (Figure 5 E,H). Both of these substrate classes are known ligands for the mammalian mannose receptor as well (Goetze et al., 2011, Kato et al., 2000). In *mrc1a* mutant situation other receptors might take over the clearance of macromolecules, which normally have lower affinity for those particular substrates and might be not as efficient in internalization of the dyes. This could possibly explain the increased number of BLECs in the *mrc1a* mutant situation: by increasing cell number and cell surface area and thereby the quantitiy of alternative scavenger receptors, mutants may attempt to compensate the loss of Mrc1a by increasing their overall capacity for the removal of accumulated waste in the brain. A candidate compensating receptor might be *mrc1b.* However, its expression has not been reported in BLECs(Venero Galanternik et al., 2017).

In addition to the common expression of *mrc1a* in BLECs and SECs, it was previously shown that BLECs also express Stabilin-2 (Bower et al., 2017), a receptor indispensable for the uptake of liposomes and hyaluronic acid from the zebrafish circulation (Campbell et al. 2018). Strikingly, and in contrast to BLECs where pHr-Avidin uptake depends on Mrc1a, pHr-Avidin is still efficiently endocytosed in *mrc1a* mutant SECs in the CV (Figure 4 A,C). This and the notion that for other substrates uptake efficiency is markedly reduced in *mrc1a* mutants, supports the conclusion that Mrc1a is important for endocytosis of many substrates within the meningeal BLECs, but is of less importance in SECs in the CV. Converseley, the analysis of *stab2* mutant fish revealed that Stabilin-2 is completely dispensable for the uptake of the endocytosis of all tested substrates in BLECs. Since Stab2 is the main receptor in liver sinusoidal endothelial cells for the binding of hyaluronic acid in mice (Schledzewski et al., 2011, Adachi and Tsujimoto, 2002), it is surprising that in *stab2* mutant fish, fluorescent hyaluronic acid can still be internalized by BLECs. This indicates, that at least in BLECs, additional receptors are involved in the endocytosis of different macromolecules. Since it has been reported that Stab1 is considered as a potential endocytosis receptor (Hansen et al., 2005) which is able to mediate the uptake of acetylated low-density protein and advanced glycation end products(Adachi and Tsujimoto, 2002), it is very likely that it plays at least a redundant role in macromolecular internalization, and this can be analyzed once the *stab1* mutants are available. Other potential receptors which would be interesting to look at are *lyve-1* and *cd44*, which have been reported to constitute hyaluronic acid receptors and to be important for dendritic cell trafficking in mammals (Johnson et al., 2017). Since *lyve-1* is also expressed in BLECs (Bower et al., 2017), it might also mediate endocytosis of hyaluronic acid in the BLECs and might play an essential role for macromolecule internalization. Yet another candidate involved in the internalization of cargo is Toll-like receptor 2 (TLR2), which forms a complex with the mannose receptor in macrophages (Tachado et al., 2007) and could be a promising candidate for taking over the endocytosis of different dyes in the *mrc1a* mutants. Notably, various TLRs including TLR2 are also expressed in human lymphatic endothelial cells (Garrafa et al., 2011). Future studies therefore need to investigate additional scavenger receptors, and double or even triple mutants might have to be employed in order to shed more light onto the molecular machinery involved in the endocytosis of proteins, liposomes, lipoproteins, polysaccharides and glycoaminoglycans.

Since the discovery of the mammalian lymphatic vasculature in the brain has recently received significant attention and has been linked to a clearance system involved in physiological and pathophysiological conditions such as ageing (Ma et al., 2017) and Alzheimer’s disease (Da Mesquita et al., 2018), the experimental analysis of BLECs provided here adds further insight into how endothelial cells contribute to the maintenance of brain homeostasis. Since BLECs have the ability to scavenge very efficiently macromolecules from the brain and qualify as scavenger endothelial cells for the brain, it raises the question which impact the absence of those cells would have on physiological and pathological conditions and whether a functionally related cell type might be conserved in mammalian brains. This needs to be addressed in future studies.

## Materials & Methods

### Zebrafish strains

Zebrafish strains were maintained under standard husbandry conditions and animal work followed guidelines of the animal ethics committees at the University of Münster, Germany. The following transgenic and mutant lines have been used in this study:

*Tg(kdr-l:HRAS-mCherry-CAAX)*^*s916*^ (Hogan et al., 2009); *Tg(lyve1:dsRed2)*^*nz101*^ (Okuda et al., 2012), *Tg(flt4:mCitrine)*^*hu7135*^ (van Impel et al., 2014), *Tg(flt1*^*enh*^:*tdTomato)*^*hu5333*^ (Bussmann et al., 2010), *Tg(mpeg1:EGFP)*^*gl22*^ (Ellett et al., 2011)

### CRISPR/Cas9

The guide RNA targeting mrc1a exon 4 (GGGGACAGTGATCCAGTGAC) was designed using chopchop algorithm (https://chopchop.cbu.uib.no/). sgRNA was synthesized as described previously (Gagnon et al., 2014). A mixture containing 15 pg of gRNA with 300pg of Cas9 mRNA were injected into the cytoplasm of one-cell stage zebrafish embryos. The −7bp deletion was identified with primers indicated in table S1.

~~~
*mrc1a* wt CTCTGGATGGGACAGTGATCCAGTGACTGGTGTATTATATCAGAGGAATGTGCAG
*mrc1a* mt CTCTGGATGGGACAGTGATC-------TGGTGTATTATATCAGAGGAATGTGCAG
~~~

### Genotyping

*mrc1a*^−*7bp*^ *and stab2* ^*– 4bp*^ embryos were genotyped by KASP using the primers indicated in Table S1.

### Injection regimes

Injections were carried out with a Pneumatic PicoPump. Embryos were anesthetized and embedded in 1.5% low melting agarose (ThermoFischer, #16520100) dissolved in embryo medium containing MS222 (Sigma, #A5040) and injected with a total volume of 0.5 nl - 1 nl per injected bolus. For intratectal injection and injection into the cerebrospinal fluid, needles were inserted into the brain in a sloped angle. Care was taken not to penetrate deep into the brain tissue.

### Dyes

The following fluorescent dyes and concentrations were used for injection: 10 kDa dextran-conjugated Alexa Fluor 647 (2mg/ml, ThermoFischer, #D22914), pHrodo Red Avidin (2 mg/ml, ThermoFischer, #P35362), pHrodo Red Dextran (2mg/mL, P10361), pHrhodo Green Dextran (2mg/mL, P35368), acLDL (1mg/ml, Thermo Fischer, L23380), oxLDL (1mg/ml, Thermo Fischer L34357), Transferrin (2mg/ml, Thermo Fischer, T23366, fluoHA Hyaluronic acid (sodium salt, 100kDa) was purchased from Lifecore Biomedical Inc. DOPG liposomes were prepared as previously described (Campbell et al. (2018).

### Imaging

Embryos were anesthetized with MS222 (Sigma, #A5040) and embedded in 1% low melting agarose (ThermoFischer, #16520100).

### Microscopy and image processing

Samples were imaged with a Leica SP8 microscope using 20x dry objectives and 40x water immersion objectives. Confocal stacks were processed using Fiji-ImageJ version 1.51g and figures were assembled using Microsoft Power Point and Adobe Photoshop and Adobe Illustrator. All data were processed using raw images with brightness, color and contrast adjusted for printing.

### Particle Analysis

Confocal maximum projections of IgG-Alexa 647 and acLDL were analyzed as follows:

~~~
roiManager(“reset”);
run(“Duplicate…”, “ “);
run(“Duplicate…”, “ “);
run(“Median…”, “radius=2”);
run(“Enhance Contrast…”, “saturated=0.6 normalize”);
run(“Threshold…”);
setThreshold(60, 255);
setOption(“BlackBackground”, true);
run(“Convert to Mask”);
run(“Analyze Particles…”, “ show=Outlines add”);
run(“Tile”);
waitForUser(“select original”);
roiManager(“deselect”);
roiManager(“multi-measure measure_all”);

roiManager(“deselect”);
~~~

Confocal maximum projections of pHR-Dextran were analyzed as follows:

~~~
roiManager(“reset”);
run(“Duplicate…”, “ “);
run(“Duplicate…”, “ “);
run(“Median…”, “radius=2”);
run(“Enhance Contrast…”, “saturated=0.4 normalize”);
run(“Threshold…”);
setThreshold(110, 255);
setOption(“BlackBackground”, true);
run(“Convert to Mask”);
run(“Analyze Particles…”, “ show=Outlines add”);
run(“Tile”);
waitForUser(“select original”);
roiManager(“deselect”);
roiManager(“multi-measure measure_all”);

roiManager(“deselect”);
~~~

### Statistical analysis

Data sets were tested for normality (Shapiro-Wilk) and equal variance. P-values were determined by Student’s t-test. When normality test failed, Mann-Whitney test was performed.

**Table S1.**
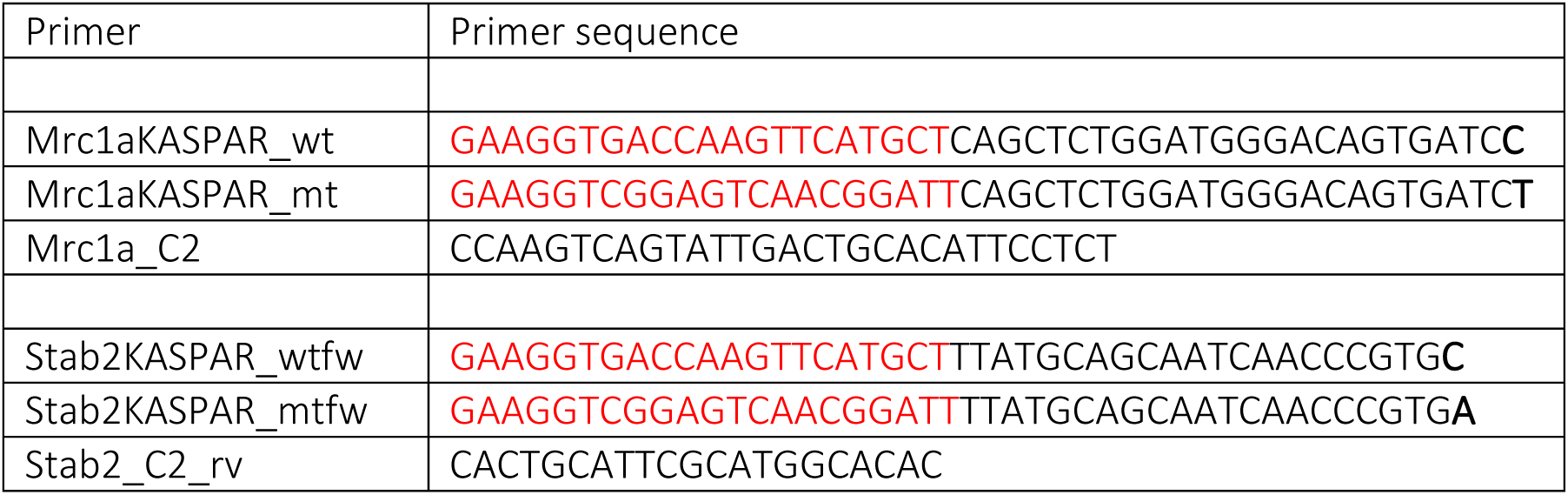

## Funding

*Deutsche Forschungsgemeinschaft (FOR2325)*

- Stefan Schulte-Merker

The funders had no role in study design, data collection and interpretation, or the decision to submit the work for publication.

## Acknowledgements

We thank members of the Schulte-Merker lab for discussions, D Stainier for providing transgenic fish lines and F. Campbell for preparing DOPG liposomes. The work was supported by the DFG (SCHU 1228/2–1, Forschergruppe FOR2325, Interactions at the Neurovascular Interface) and the CiM Cluster of Excellence (EXC 1003 CiM, WWU Münster, Germany). Furthermore we thank Thomas Zobel for programming the macro for the quantification of the fluorescence intesity in Fiji.

## Ethics

Animal experimentation: Experimental procedures were conducted under project licence.

---

**Supplemental Figure 1:**
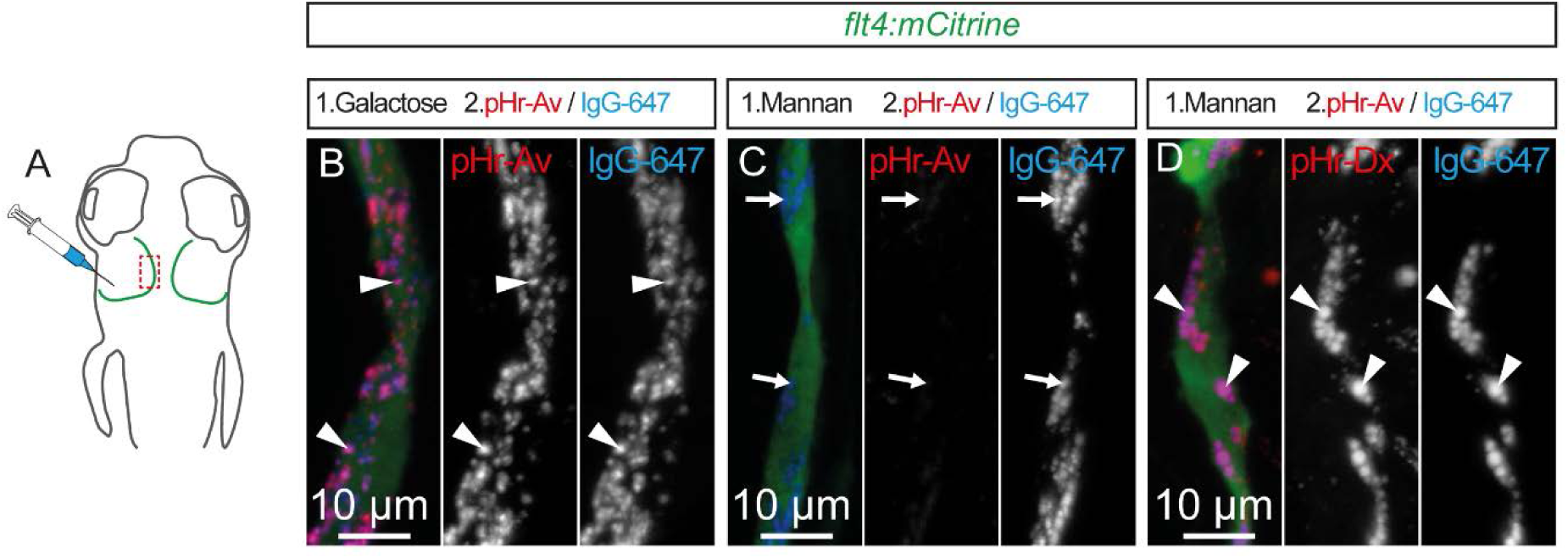
Separate, consecutive injection of fluorescent dyes over time labels the same lyso-endosomal compartments. (A) Overview of the zebrafish head region depicting intratectal injection of compounds and fluorescent dyes into the center of the TeO close to the meninges in 5dpf embryos. Red inset denotes area of image detail for representative dorsal confocal projections of BLECs in B-D. (B – D) Numbers in headers denote order of separate, consecutive injections. Consecutive injection of indicated fluorescent dyes and mannan (separated by a ten minutes time interval) over time labels the same lyso-endosomal compartments. After initial administration and uptake, the injection of mannan blocks further uptake of pHr-Av. At 20 minutes after the administration of IgG-647, most pHr-Av positive lyso-endosomal compartments are IgG-647 negative. After further incubation, at 60 minutes, the majority of pHr-Av positive compartments show accumulation of IgG-647 tracer. BLEC, brain lymphatic endothelial cell; dpf, days post fertilization; IgG-647, IgG-conjugated Alexa Fluor 674; pHr-Av, pHrodo™ Red Avidin; TeO, Optic Tectum.

**Supplemental Figure 2:**
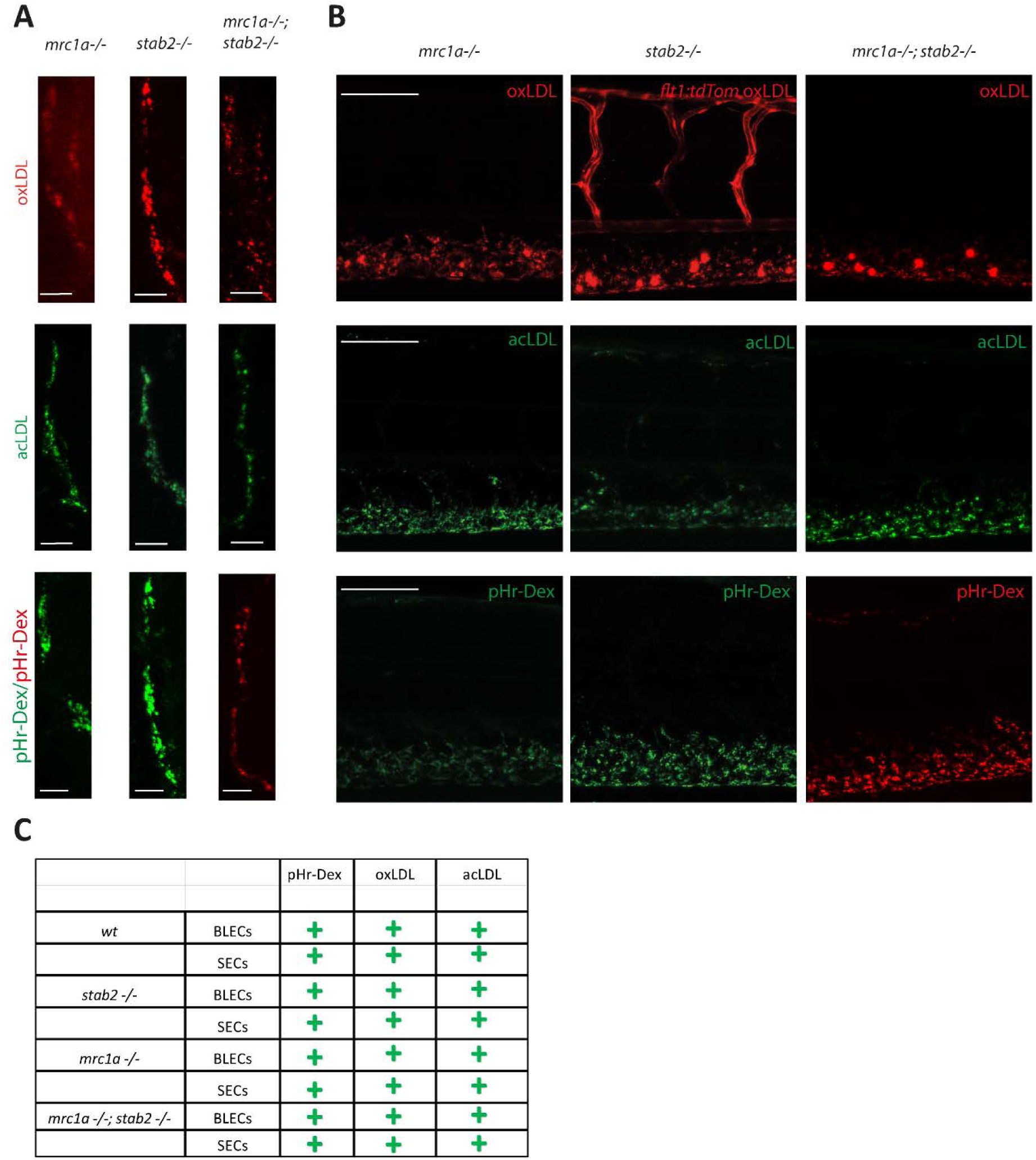
Involvement of Stab2 and Mrc1a receptor during cargo uptake by BLECs and SECs. Analysis of dye uptake of SECs and BLECs of either *mrc1a* ^−*/*−^ or *stab2*^−*/*−^ single mutants, or of *mrc1a*^−*/*−^; *stab2*^−*/*−^ double mutant embryos. (A+B) Confocal images of cargo injections showing the accumulation of oxLDL and acLDL phR-Dextan in BLECs after injection into the CSF or the optic tectum (A) and SECs of the CV after dye administration into the bloodstream of embryos (B) with the indicated genotypes. (C) Summary of the fluorescent substrate molecule uptake experiments. pHr-Dex – pHr-Dextrane; acLDL-acetylated LDL; oxLDL-oxidated LDL

## Literature

Absinta, M., Ha, S. K., Nair, G., Sati, P., Luciano, N. J., Palisoc, M., Louveau, A., Zaghloul, K. A., Pittaluga, S., Kipnis, J. & Reich, D. S. 2017. Human and nonhuman primate meninges harbor lymphatic vessels that can be visualized noninvasively by MRI. Elife, 6.

Adachi, H. & Tsujimoto, M. 2002. FEEL-1, a novel scavenger receptor with in vitro bacteria-binding and angiogenesis-modulating activities. J Biol Chem, 277, 34264–70.

Antila, S., Karaman, S., Nurmi, H., Airavaara, M., Voutilainen, M. H., Mathivet, T., Chilov, D., Li, Z., Koppinen, T., Park, J. H., Fang, S., Aspelund, A., Saarma, M., Eichmann, A., Thomas, J. L. & Alitalo, K. 2017. Development and plasticity of meningeal lymphatic vessels. J Exp Med, 214, 3645–3667.

Aspelund, A., Antila, S., Proulx, S. T., Karlsen, T. V., Karaman, S., Detmar, M., Wiig, H. & Alitalo, K. 2015. A dural lymphatic vascular system that drains brain interstitial fluid and macromolecules. J Exp Med, 212, 991–9.

Barron, K. D. 1995. The microglial cell. A historical review. J Neurol Sci, 134 Suppl, 57–68.

Bower, N. I., Koltowska, K., Pichol-Thievend, C., Virshup, I., Paterson, S., Lagendijk, A. K., Wang, W., Lindsey, B. W., Bent, S. J., Baek, S., Rondon-Galeano, M., Hurley, D. G., Mochizuki, N., Simons, C., Francois, M., Wells, C. A., Kaslin, J. & Hogan, B. M. 2017. Mural lymphatic endothelial cells regulate meningeal angiogenesis in the zebrafish. Nat Neurosci, 20, 774–783.

Bruch, R. C. & White, H. B. 1982. Compositional and structural heterogeneity of avidin glycopeptides. Biochemistry, 21, 5334–5341.

Bussmann, J., Bos, F. L., Urasaki, A., Kawakami, K., Duckers, H. J. & Schulte-Merker, S. 2010. Arteries provide essential guidance cues for lymphatic endothelial cells in the zebrafish trunk. Development, 137, 2653–7.

Campbell, F., Bos, F. L., Sieber, S., Arias-Alpizar, G., Koch, B. E., Huwyler, J., Kros, A. & Bussmann, J. 2018. Directing Nanoparticle Biodistribution through Evasion and Exploitation of Stab2-Dependent Nanoparticle Uptake. ACS Nano, 12, 2138–2150.

Chen, J., He, J., Ni, R., Yang, Q., Zhang, Y. & Luo, L. 2019. Cerebrovascular Injuries Induce Lymphatic Invasion into Brain Parenchyma to Guide Vascular Regeneration in Zebrafish. Dev Cell, 49, 697–710.e5.

Da Mesquita, S., Louveau, A., Vaccari, A., Smirnov, I., Cornelison, R. C., Kingsmore, K. M., Contarino, C., Onengut-Gumuscu, S., Farber, E., Raper, D., Viar, K. E., Powell, R. D., Baker, W., Dabhi, N., Bai, R., Cao, R., Hu, S., Rich, S. S., Munson, J. M., Lopes, M. B., Overall, C. C., Acton, S. T. & Kipnis, J. 2018. Functional aspects of meningeal lymphatics in ageing and Alzheimer’s disease. Nature, 560, 185–191.

Delange, R. J. 1970. Egg White Avidin: I. AMINO ACID COMPOSITION; SEQUENCE OF THE AMINO-AND CARBOXYL-TERMINAL CYANOGEN BROMIDE PEPTIDES. Journal of Biological Chemistry, 245, 907–916.

Ellett, F., Pase, L., Hayman, J. W., Andrianopoulos, A. & Lieschke, G. J. 2011. mpeg1 promoter transgenes direct macrophage-lineage expression in zebrafish. Blood, 117, e49–56.

Fiete, D., Beranek, M. C. & Baenziger, J. U. 1997. The macrophage/endothelial cell mannose receptor cDNA encodes a protein that binds oligosaccharides terminating with SO_4_-4-GalNAcβ1,4GlcNAcβ or Man at independent sites. Proceedings of the National Academy of Sciences, 94, 11256–11261.

Gagnon, J. A., Valen, E., Thyme, S. B., Huang, P., Akhmetova, L., Pauli, A., Montague, T. G., Zimmerman, S., Richter, C. & Schier, A. F. 2014. Efficient mutagenesis by Cas9 protein-mediated oligonucleotide insertion and large-scale assessment of single-guide RNAs. PLoS One, 9, e98186.

Garrafa, E., Imberti, L., Tiberio, G., Prandini, A., Giulini, S. M. & Caimi, L. 2011. Heterogeneous expression of toll-like receptors in lymphatic endothelial cells derived from different tissues. Immunol Cell Biol, 89, 475–81.

Goetze, A. M., Liu, Y. D., Zhang, Z., Shah, B., Lee, E., Bondarenko, P. V. & Flynn, G. C. 2011. High-mannose glycans on the Fc region of therapeutic IgG antibodies increase serum clearance in humans. Glycobiology, 21, 949–959.

Goldmann, T., Wieghofer, P., Jordao, M. J., Prutek, F., Hagemeyer, N., Frenzel, K., Amann, L., Staszewski, O., Kierdorf, K., Krueger, M., Locatelli, G., Hochgerner, H., Zeiser, R., Epelman, S., Geissmann, F., Priller, J., Rossi, F. M., Bechmann, I., Kerschensteiner, M., Linnarsson, S., Jung, S. & Prinz, M. 2016. Origin, fate and dynamics of macrophages at central nervous system interfaces. Nat Immunol, 17, 797–805.

Green, N. M. & Toms, E. J. 1970. Purification and crystallization of avidin. Biochemical Journal, 118, 67–70.

Hansen, B., Longati, P., Elvevold, K., Nedredal, G. I., Schledzewski, K., Olsen, R., Falkowski, M., Kzhyshkowska, J., Carlsson, F., Johansson, S., Smedsrod, B., Goerdt, S., Johansson, S. & Mccourt, P. 2005. Stabilin-1 and stabilin-2 are both directed into the early endocytic pathway in hepatic sinusoidal endothelium via interactions with clathrin/AP-2, independent of ligand binding. Exp Cell Res, 303, 160–73.

Hogan, B. M., Bos, F. L., Bussmann, J., Witte, M., Chi, N. C., Duckers, H. J. & Schulte-Merker, S. 2009. Ccbe1 is required for embryonic lymphangiogenesis and venous sprouting. Nat Genet, 41, 396–8.

Hubbard, A. L., Wilson, G., Ashwell, G. & Stukenbrok, H. 1979. An electron microscope autoradiographic study of the carbohydrate recognition systems in rat liver. I. Distribution of 125I-ligands among the liver cell types. The Journal of Cell Biology, 83, 47–64.

Johnson, L. A., Banerji, S., Lawrance, W., Gileadi, U., Prota, G., Holder, K. A., Roshorm, Y. M., Hanke, T., Cerundolo, V., Gale, N. W. & Jackson, D. G. 2017. Dendritic cells enter lymph vessels by hyaluronan-mediated docking to the endothelial receptor LYVE-1. Nature Immunology, 18, 762.

Kato, M., Neil, T. K., Fearnley, D. B., Mclellan, A. D., Vuckovic, S. & Hart, D. N. J. 2000. Expression of multilectin receptors and comparative FITC–dextran uptake by human dendritic cells. International Immunology, 12, 1511–1519.

Kettenmann, H. 2007. The brain’s garbage men. Nature, 446, 987.

Lee, S. J., Evers, S., Roeder, D., Parlow, A. F., Risteli, J., Risteli, L., Lee, Y. C., Feizi, T., Langen, H. & Nussenzweig, M. C. 2002. Mannose receptor-mediated regulation of serum glycoprotein homeostasis. Science, 295, 1898–901.

Louveau, A., Smirnov, I., Keyes, T. J., Eccles, J. D., Rouhani, S. J., Peske, J. D., Derecki, N. C., Castle, D., Mandell, J. W., Lee, K. S., Harris, T. H. & Kipnis, J. 2015. Structural and functional features of central nervous system lymphatic vessels. Nature, 523, 337–41.

Ma, Q., Ineichen, B. V., Detmar, M. & Proulx, S. T. 2017. Outflow of cerebrospinal fluid is predominantly through lymphatic vessels and is reduced in aged mice. Nature Communications, 8, 1434.

Malovic, I., SØRensen, K. K., Elvevold, K. H., Nedredal, G. I., Paulsen, S., Erofeev, A. V., SmedsrØD, B. H. & Mccourt, P. A. G. 2007. The mannose receptor on murine liver sinusoidal endothelial cells is the main denatured collagen clearance receptor. Hepatology, 45, 1454–1461.

Mascagni, P. & Bellini, G. B. 1816. Istoria completa dei vasi linfatici, E. Pacini.

Mazaheri, F., Breus, O., Durdu, S., Haas, P., Wittbrodt, J., Gilmour, D. & Peri, F. 2014. Distinct roles for BAI1 and TIM-4 in the engulfment of dying neurons by microglia. Nature Communications, 5, 4046.

Metcalf, D. J., Garcia-Arencibia, M., Hochfeld, W. E. & Rubinsztein, D. C. 2012. Autophagy and misfolded proteins in neurodegeneration. Exp Neurol, 238, 22–8.

Mi, Y., Shapiro, S. D. & Baenziger, J. U. 2002. Regulation of lutropin circulatory half-life by the mannose/N-acetylgalactosamine-4-SO4 receptor is critical for implantation in vivo. J Clin Invest, 109, 269–76.

Okuda, K. S., Astin, J. W., Misa, J. P., Flores, M. V., Crosier, K. E. & Crosier, P. S. 2012. lyve1 expression reveals novel lymphatic vessels and new mechanisms for lymphatic vessel development in zebrafish. Development, 139, 2381–91.

Platt, N., Da Silva, R. P. & Gordon, S. 1998. Recognizing death: the phagocytosis of apoptotic cells. Trends Cell Biol, 8, 365–72.

Sallusto, F., Cella, M., Danieli, C. & Lanzavecchia, A. 1995. Dendritic cells use macropinocytosis and the mannose receptor to concentrate macromolecules in the major histocompatibility complex class II compartment: downregulation by cytokines and bacterial products. J Exp Med, 182, 389–400.

Schledzewski, K., Geraud, C., Arnold, B., Wang, S., Grone, H. J., Kempf, T., Wollert, K. C., Straub, B. K., Schirmacher, P., Demory, A., Schonhaber, H., Gratchev, A., Dietz, L., Thierse, H. J., Kzhyshkowska, J. & Goerdt, S. 2011. Deficiency of liver sinusoidal scavenger receptors stabilin-1 and -2 in mice causes glomerulofibrotic nephropathy via impaired hepatic clearance of noxious blood factors. J Clin Invest, 121, 703–14.

Schulte-Merker, S., Sabine, A. & Petrova, T. V. 2011. Lymphatic vascular morphogenesis in development, physiology, and disease. J Cell Biol, 193, 607–18.

Seternes, T., Sorensen, K. & Smedsrod, B. 2002. Scavenger endothelial cells of vertebrates: a nonperipheral leukocyte system for high-capacity elimination of waste macromolecules. Proc Natl Acad Sci U S A, 99, 7594–7.

Tachado, S. D., Zhang, J., Zhu, J., Patel, N., Cushion, M. & Koziel, H. 2007. Pneumocystis-mediated IL-8 release by macrophages requires coexpression of mannose receptors and TLR2. J Leukoc Biol, 81, 205–11.

Van Impel, A., Zhao, Z., Hermkens, D. M., Roukens, M. G., Fischer, J. C., Peterson-Maduro, J., Duckers, H., Ober, E. A., Ingham, P. W. & Schulte-Merker, S. 2014. Divergence of zebrafish and mouse lymphatic cell fate specification pathways. Development, 141, 1228–38.

Van Lessen, M., Shibata-Germanos, S., Van Impel, A., Hawkins, T. A., Rihel, J. & Schulte-Merker, S. 2017. Intracellular uptake of macromolecules by brain lymphatic endothelial cells during zebrafish embryonic development. Elife, 6.

Venero Galanternik, M., Castranova, D., Gore, A. V., Blewett, N. H., Jung, H. M., Stratman, A. N., Kirby, M. R., Iben, J., Miller, M. F., Kawakami, K., Maraia, R. J. & Weinstein, B. M. 2017. A novel perivascular cell population in the zebrafish brain. Elife, 6.

